# NOSA, an analytical toolbox for multicellular optical electrophysiology

**DOI:** 10.1101/2020.03.15.992420

**Authors:** Sebastian Oltmanns, Frauke Sophie Abben, Anatoli Ender, Sophie Aimon, Richard Kovacs, Stephan J Sigrist, Douglas A Storace, Jörg R P Geiger, Davide Raccuglia

**Affiliations:** Institute of Neurophysiology, Charité – Universitätsmedizin Berlin, corporate member of Freie Universität Berlin and Humboldt-Universität zu Berlin, and Berlin Institute of Health, Charitéplatz 1, 10117 Berlin, Germany; German Center for Neurodegenerative Disorders, Charité – Universitätsmedizin Berlin, Charitéplatz 1, 10117 Berlin, Germany; School of Life Sciences, Technical University Munich, 85354 Freising, Germany; Institute for Biology/Genetics, Freie Universität Berlin, Takustrasse 6, 14195 Berlin, Germany; NeuroCure, Charité – Universitätsmedizin Berlin, corporate member of Freie Universität Berlin and Humboldt-Universität zu Berlin, and Berlin Institute of Health, Charitéplatz 1, 10117 Berlin, Germany; Florida State University, Department of Biological Science, Tallahassee, FL 32306

## Abstract

Understanding how neural networks generate activity patterns and communicate with each other requires monitoring the electrical activity from many neurons simultaneously. Perfectly suited tools for addressing this challenge are genetically encoded voltage indicators (GEVIs) because they can be targeted to specific cell types and optically report the electrical activity of individual, or populations of neurons. However, analyzing and interpreting the data from voltage imaging experiments is challenging because high recording speeds and properties of current GEVIs yield only low signal-to-noise ratios, making it necessary to apply specific analytical tools. Here, we present NOSA (Neuro-Optical Signal Analysis), a novel open source software designed for analyzing voltage imaging data and identifying temporal interactions between electrical activity patterns of different origin.

In this manuscript we explain the challenges that arise during voltage imaging experiments and provide hands-on analytical solutions. We demonstrate how NOSA’s baseline fitting, filtering algorithms and movement correction can compensate for shifts in baseline fluorescence and extract electrical patterns from low signal-to-noise recordings. Moreover, NOSA contains powerful features to identify oscillatory frequencies in electrical patterns and extract neuronal firing characteristics. NOSA is the first open-access software to provide an option for analyzing simultaneously recorded optical and electrical data derived from patch-clamp or other electrode-based recordings. To identify temporal relations between electrical activity patterns we implemented different options to perform cross correlation analysis, demonstrating their utility during voltage imaging in *Drosophila* and mice. All features combined, NOSA will facilitate the first steps into using GEVIs and help to realize their full potential for revealing cell-type specific connectivity and functional interactions. If you would like to test NOSA, please send an email to the lead contact.

## Introduction

One goal of The American BRAIN initiative was to develop methods to comprehend complex activity patterns in specific brain networks and even in whole brains (Alivisatos et al., 2012). One crucial step towards gaining insight into the mechanisms and interactions of neural activity patterns is the development of tools that allow for measuring multi-cellular electrical activity and analyzing their complex datasets (Alivisatos et al., 2012).

Genetically encoded voltage indicators (GEVIs) have emerged as promising tools for measuring neural electrical activity (Lin and Schnitzer, 2016; Yang and St-Pierre, 2016). GEVIs are powerful tools, in part because they can be genetically targeted to specific neural populations and optically report the electrical activity from many neurons simultaneously, and even from neuropil that is otherwise inaccessible to classical electrophysiology. GEVIs have been successfully used for monitoring multicellular activity and population dynamics in *Drosophila* (Cao et al., 2013; Raccuglia et al., 2016; Aimon et al., 2019; Raccuglia et al., 2019), visual and olfactory responses in mice (Gong et al., 2015; Storace et al., 2015; Storace and Cohen, 2017), cerebellar activity in Zebrafish (Miyazawa et al., 2018) and also pharyngeal activity in *C. elegans* (Azimi Hashemi et al., 2019).

Although GEVIs are being continually improved (Lin and Schnitzer, 2016; Storace et al., 2016), high recording speeds, low signal-to-noise ratios (SNR) and GEVI-specific kinetics bring about unique challenges with respect to data analysis (Yang and St-Pierre, 2016; Kulkarni and Miller, 2017) and thus require the development of adequate processing tools. Yet, there is currently no freely available software, which combines processing tools addressing these challenges with analytical tools for identifying specific activity patterns, temporal relations and functional interactions.

Here we present NOSA (Neuro-Optical Signal Analysis) – an open source software designed specifically for the analysis of multicellular optical electrophysiology. NOSA features baseline fitting and filtering algorithms to extract electrical patterns from high speed recordings with low SNR. Moreover, NOSA provides analytical tools for identifying specific activity patterns and their temporal relation via functions that provide spectral and cross-correlation analysis. NOSA also includes features for spike- and burst detection, movement artefact compensation, and the ability to analyze simultaneously performed optical and electrical recordings. With these analytical tools, intuitive design, and convenient graphical interface, NOSA should greatly facilitate the first steps into using GEVIs, enabling laboratories around the world to perform and analyze multicellular voltage imaging recordings.

## Results

### NOSA Interface and overview of processing tools

Our software package NOSA (**Fig. 1**) is designed to process and analyze voltage imaging recordings. Recordings can be imported into NOSA as tif/tiff files, which can be temporally cropped (the user can select a specific time window) or corrected for movement artefacts (**Fig. 1A**). NOSA automatically calculates the relative changes in fluorescence for selected regions of interest (ROIs) (**Fig. 1B, C**) after factoring in recording speed, background correction as well as selected fitting (e.g., exponential drift correction) and filtering algorithms (**Fig. 1D**). Activity patterns from different ROIs (e.g., cells) can be displayed in one plot to facilitate comparisons, although the user can easily switch to a more detailed view of the currently selected ROI (**Fig. 1E**).

**Figure 1.**
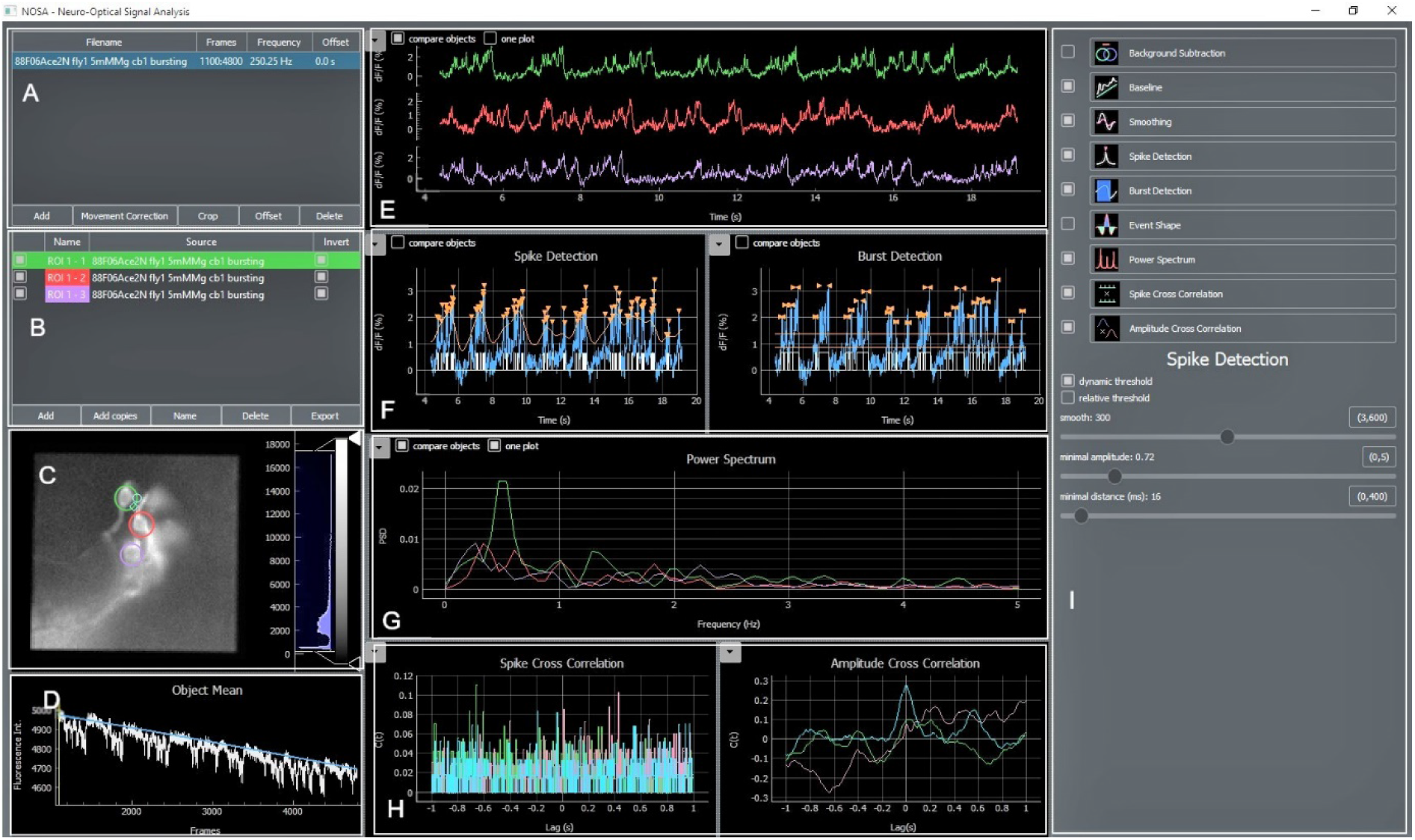
NOSA Interface. **A,** The file manager shows imported files and contains options for movement correction, temporal cropping, offset correction and interpolation of the recording speed. **B,** ROI manager showing added ROIs, which can be renamed and deleted. ROI masks can be generated by copying ROIs into other files. **C**, Video panel showing currently selected frame and added ROIs. The recording can be adjusted for brightness and contrast. **D,** Average fluorescence intensity of selected ROIs with baseline fitting curve and slide for selecting specific frames of the recording. **E**, Processed data showing relative changes in fluorescence over time. Calculation of the relative change in fluorescence is based on recording speed, background subtraction, baseline correction and smoothing. **F**, Panel for semi-automatic detection of spikes and bursts. Detected events are indicated with orange arrows and white lines/squares indicating spikes/bursts. **G**, Power spectrum analysis based on processed data. If the event shape feature is selected this panel also shows the average shape of detected events. **H**, Panel showing spike cross correlation calculated based on detected spikes and amplitude cross correlation calculated based on relative changes in fluorescence. **I,** Control panel for adjusting the settings of each feature.

Because most GEVIs exhibit decreases in their fluorescence in response to depolarization, NOSA provides the option to invert the relative changes (**Fig. 1B**). To generate ROI masks, selected ROIs can be copied and pasted into other recordings (**Fig. 1B**). All data extracted by NOSA can be exported as spreadsheet files (**Fig. 1B**).

As high recording speeds generate a larger number of frames, we included simple but efficient options to increase software performance. For example, the recording frequency can be reduced by applying different interpolation algorithms (**Fig. 1B,** right-click on file name). To simplify the simultaneous evaluation of multiple recordings this method can also be used to unify different recording speeds. However, this method should only be used when the recording frequency is higher than needed to resolve single depolarization events. Moreover, software performance can be increased by using a rectangular ROI instead of an elliptical and by deselecting the live preview mode. In the live preview mode, changes in relative fluorescence are displayed while moving around a ROI. After deselecting live preview mode (**Fig. 1C,** right-click), changes in relative fluorescence are only displayed after placing the ROI.

NOSA features power spectrum analysis as well as spike and burst detection to analyze activity patterns (**Fig. 1F, G**). The event shape feature uses detected events to display the average firing characteristics of a neuron. To analyze temporal relations and functional interactions between activity patterns, cross correlation can be performed on detected events and on the relative changes in fluorescence (**Fig. 1H**). Via the control panel all features can be controlled (**Fig. 1I**) for each ROI independently and settings selected for one ROI can easily be applied to all other ROIs.

### Multicellular optical electrophysiology

To demonstrate the utility of NOSA we used *Drosophila* to express the genetically encoded voltage indicator (GEVI) ArcLight in ellipsoid body R5 ring neurons and the GEVI Varnam in fan-shaped neurons (**Fig. 2**). Both neural structures are considered to be integration centers for various sensory modalities (Seelig and Jayaraman, 2013; Green et al., 2017; Sun et al., 2017; Hu et al., 2018) and play important roles in locomotion (Strauss and Heisenberg, 1993) and sleep regulation (Donlea et al., 2014; Liu et al., 2016; Donlea et al., 2018; Guo et al., 2018; Raccuglia et al., 2019).

**Figure 2.**
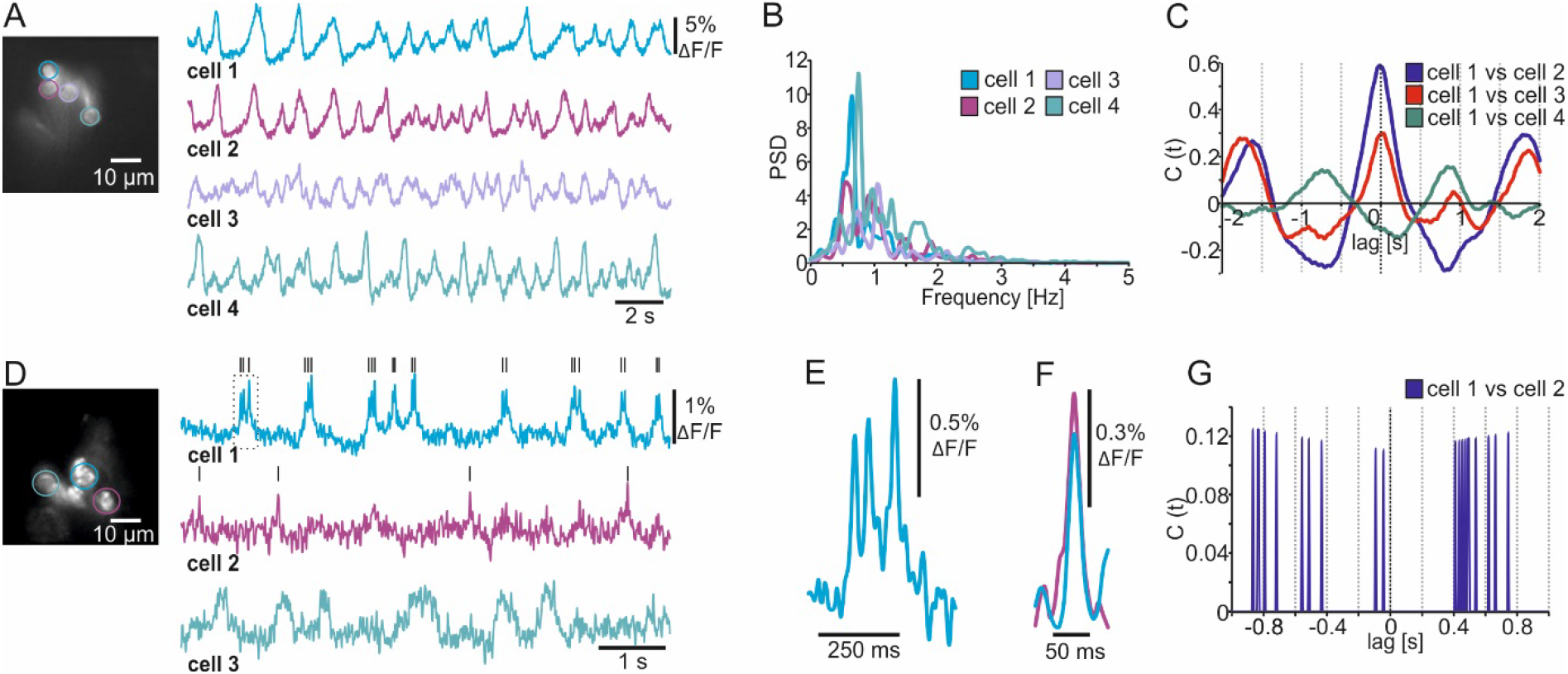
Analyzing multicellular electrical activity from optical recordings. **A,** Wide-field image and single-cell membrane potential oscillations of *Drosophila* R5 ring neurons expressing the GEVI ArcLight. **B,** Power spectrum analysis of R5 electrical activity shown in A. **C,** Amplitude cross correlation indicating the temporal relation of R5 electrical patterns shown in A. **D,** Wide-field image and single-cell electrical activity of fan-shaped body neurons expressing the GEVI Varnam. Lines indicate detected spikes using a linear threshold approach. **E,** Burst (see dashed square in D) showing that Varnam resolves single spikes during bursting. **F,** The event shape feature shows the average kinetics of the action potentials detected in cell 1 and 2 (see black lines in D). **G,** Spike cross correlation indicating no temporal relations between spikes detected in D.

After setting a ROI, the optical trace should first be corrected for shifts in baseline fluorescence (e.g. due to photoisomerization and bleaching). Several baseline correction algorithms are included (**Suppl. Fig. 1-3**), and the user can set additional baseline markers to guide the fitting curve for more complex shifts in baseline fluorescence (**Suppl. Fig. 1A**). After baseline correction, smoothing and inversion of the raw fluorescence (depolarization leads to a reduction in fluorescence), the electrical patterns of single R5 ring neurons become apparent (**Fig. 2A, compare Suppl. Fig. 1**). A power spectrum analysis revealed that single R5 neurons oscillate between 0.5 – 1.5 Hz (**Fig. 2B**). We recently reported that R5 oscillations within this spectrum are linked to the fly’s sleep quality because they facilitate consolidated sleep phases (Raccuglia et al., 2019). The cross-correlation function built into NOSA provides a simple way of visualizing the temporal relation between the electrical patterns of the different cells (**Fig. 2C**). The cross correlogram indicates that electrical patterns of cells 1, 2 and 3 largely overlap (main phase lag at 0) while cell 4 is out of phase (**Fig. 2C**). We also implemented the option to perform a cross correlation on instantaneous amplitudes (Adhikari et al., 2010) and use a band pass filter for comparing temporal relations within a specific frequency range.

Due to the relatively low recording speed (78 Hz) and the slow kinetics of ArcLight we could not resolve single spikes in this example (**Fig. 2A**). By increasing the recording speed to 160 Hz and taking advantage of the improved kinetics of the red-shifted GEVI Varnam (Kannan et al., 2018), we resolved single action potentials within the electrical activity of dorsal fan-shaped body neurons (**Fig. 2D, E**). Compared to ArcLight, the SNR is lower and thus the detection of spikes heavily depends on the imaging conditions (**Fig. 2D**), as individual spikes could not be resolved in the dimmer cells (**Fig. 2D, see cell3**).

That said, several functions are included to facilitate spike detection in low signal-to-noise recordings. This includes several filtering algorithms (see next chapter) as well as the ability to semi-automatically detect spikes and visualize their average shape using the event-shape feature (**Fig. 2F, compare Suppl. Fig. 2**). Moreover, the temporal relation between detected spikes can be analyzed using the spike cross correlation feature (**Fig. 2G, compare Suppl. Fig. 2**).

### Spike and burst detection

In NOSA, electrical characteristics of single neurons as well as the kinetics of different GEVIs can be analyzed in detail using event detection features. To demonstrate this, we compare the GEVIs ArcLight and Ace2N (Gong et al., 2015) in R5 neurons in *Drosophila*. While most R5 neurons burst (~90%), some mainly spike (Liu et al., 2016). NOSA’s spike detection and event shape feature was used to analyze two spiking R5 neurons expressing either ArcLight or Ace2N (**Fig. 3A-B**). While the kinetics of the depolarization are comparable, the repolarization is considerably slower in ArcLight, which is in accordance with previous findings indicating that Ace2N has faster kinetics (Gong et al., 2015).

**Figure 3.**
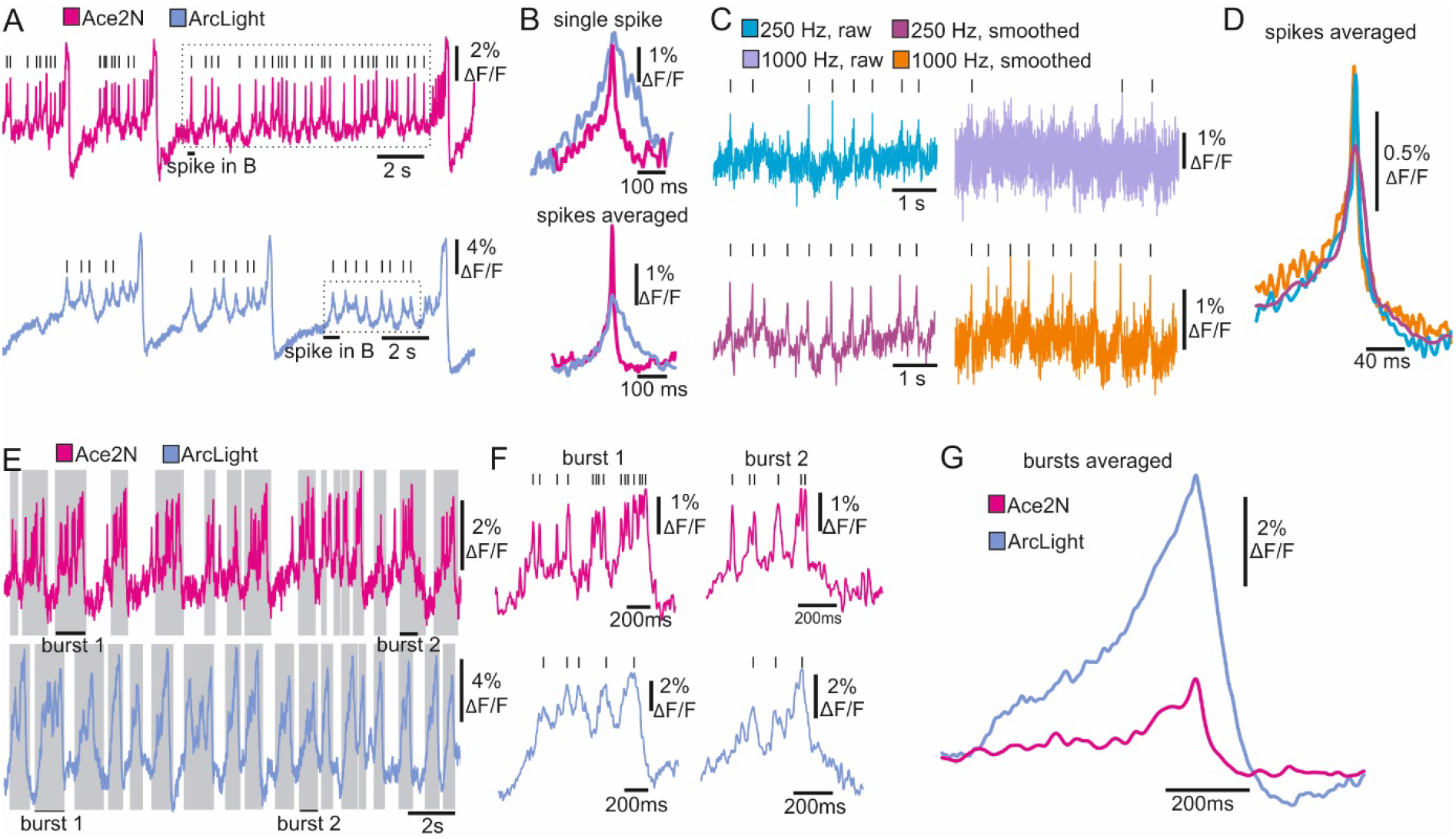
Spike and burst detection revealing kinetic properties of GEVIs and neuron-specific firing characteristics. **A,** Optical recordings of spiking R5 neurons expressing the GEVIs Ace2N and ArcLight. Lines indicate detected spikes using a linear threshold approach. **B,** Single and averaged spikes selected from recordings in A (dashed square indicates spikes selected for event shape). The event shape feature indicates improved kinetics of Ace2N. **C,** Optical recordings of a spiking R5 neuron (Ace2N) performed at different recording speeds. Lines indicate identified spikes. **D,** The event shape feature shows the average kinetics of spikes detected in C. **E, F,** Optical recordings of bursting R5 neurons expressing the GEVIs Ace2N and ArcLight. Grey shaded areas and lines indicate detected bursts and spikes using the dynamic threshold approach. **G,** The event shape feature generates the averaged kinetics of detected bursts indicating GEVI-specific differences in the optical representation of bursts.

Optical representations of action potentials do not only depend on the GEVI kinetics, but also on a sensitive interplay between recording-speed and signal-to-noise ratio. To demonstrate this, action potentials were imaged using Ace2N at either 250 Hz or 1000 Hz (**Fig. 3C**). The faster recording speed resulted in a drastically reduced signal-to-noise ratio, making action potentials barely detectable (**Fig. 3C**). However, filtering algorithms provided in NOSA can increase the signal-to-noise ratio and thus the spike detection fidelity, as demonstrated here by using the Savitzky-Golay algorithm (**Fig. 3C**). Automatic averaging of detected spikes (event shape) indicates that the temporal features of single action potentials are well represented at 250 Hz (**Fig. 3D**). Here, the limiting factors seem to be the temporal dynamics of the GEVI itself.

An advantage of enhanced temporal dynamics of a GEVI becomes apparent when analyzing high-frequency spikes in bursting neurons (**Fig. 3E**). Using Ace2N, spikes riding on top of bursts are more likely to be resolved and are therefore more readily detectable (**Fig. 3F**). However, the event shape feature indicates that the temporal characteristics of the bursts are identical but ArcLight produces a bigger change in relative fluorescence (**Fig. 3G**). This is likely due to the slower kinetics of ArcLight, resulting in several spikes probably merging into one “spike”.

### Movement and background correction

Due to a relatively small SNR, movement artefacts are especially problematic for *in vivo* voltage imaging. We therefore implemented several movement correction algorithms into NOSA. We here demonstrate the symmetric diffeomorphic algorithm, which was originally designed for detecting brain deformations during magnetic resonance imaging (Avants et al., 2008). This sophisticated algorithm is very time consuming but shows impressive results (**Fig. 4A**). Single R5 neurons recorded *in vivo* show substantial movement artefacts (**Fig. 4A**). However, after performing the symmetric diffeomorphic algorithm electrical activity can be recovered even during periods of heavy movement.

**Figure 4.**
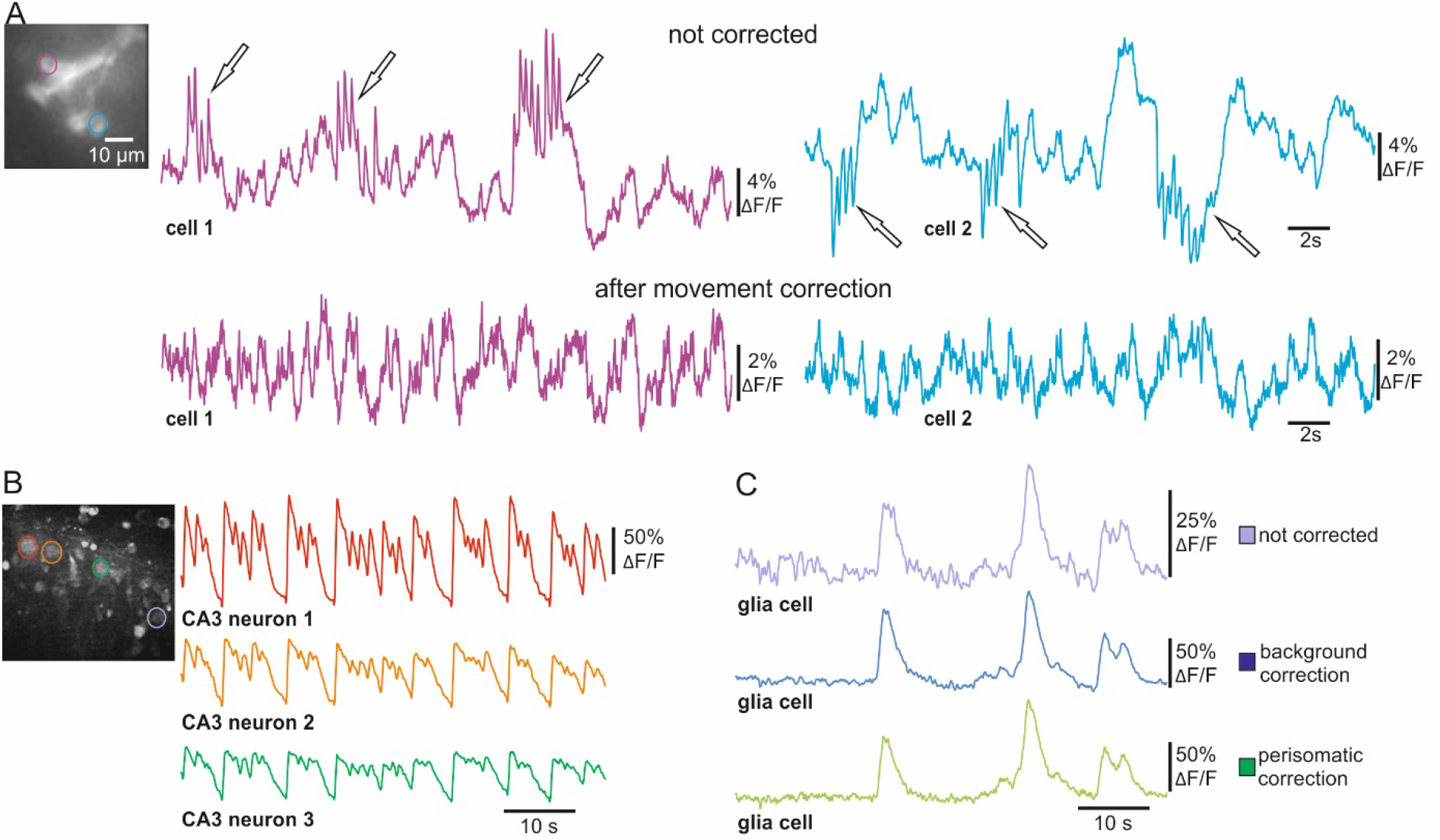
Movement and background correction for voltage and Ca^2+^ imaging. **A,** *In vivo* voltage imaging (ArcLight) of single R5 neurons before and after performing the symmetric diffeomorphic movement correction. **B,** Ca^2+^ imaging from hippocampal slices shows synchronous activity of CA3 neurons after inducing epileptiform activity. **C,** Activity of glia cell indicated in wide-field image in B before and after ROI background correction and perisomatic background correction.

Scattering light can also reduce the signal-to-noise ratio in epifluorescence imaging experiments. This is especially problematic in bulk dye-loading procedures as Ca^2+^- or voltage sensitive dyes accumulate differently in different cell-types (i.e. neurons and glial cells), leading to a large variability in fluorescence intensity. To address this issue, we implemented ROI and perisomatic background correction algorithms into NOSA (**Fig. 4B, C**). To demonstrate their utility, we analyzed epileptiform activity in hippocampal slices (Kovacs et al., 2001) (**Fig. 4B**). Here, strong and synchronized increases in fluorescence in CA3 neurons lead to increased light-scattering, affecting the SNR in glial cells. However, ROI and perisomatic background correction both successfully reduce the effects of scattered light, increasing the SNR in the activity pattern of a glial cell (**Fig. 4C**).

### Combined *in vivo* optical and classical electrophysiology

Being able to compare optical measurements alongside ongoing electrical recordings is useful in many ways. To our knowledge, NOSA is the first open-access tool to provide the option for analyzing optical and electrical traces in parallel. For example, we performed simultaneous patch-clamp and optical recordings *in vivo* from R5 ring neurons and uploaded optical and electrical traces (abf files/ axon binary file format) into NOSA (**Suppl. Fig. 3**). As these traces may be recorded with different systems, temporal delays between the systems could falsify a direct comparison. We implemented an offset function into NOSA, which allows shifting one trace in relation to the other (**Suppl. Fig. 3**). Comparing optical and electrical traces demonstrates that ArcLight faithfully represents changes in membrane potential (**Fig. 5A-C**) and that a change in relative fluorescence of 2% approximates a change of 23 mV in membrane potential (**Fig. 5A-C**). However, the relation between changes in fluorescence and absolute membrane potential will highly depend on the expression strength and must therefore be determined for each cell type. To simplify a direct comparison NOSA can adjust the sampling rate via interpolation of data points. In this example we reduced the sampling rate of the electrical trace from 10 kHz to 2 kHz smoothing out the spikes on top of the bursts which in this case are not represented in the optical trace (**Fig. 5A**).

**Figure 5.**
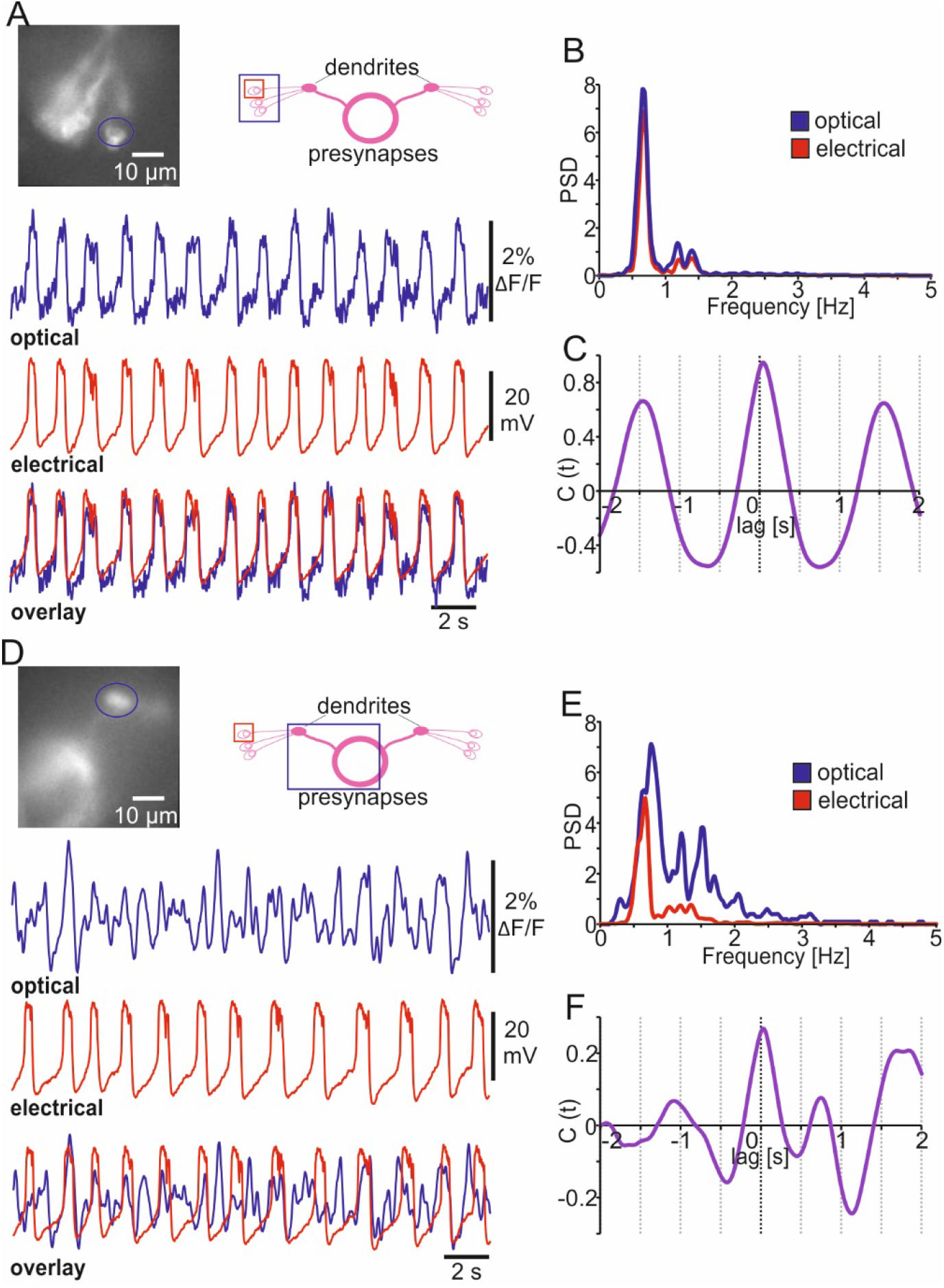
Simultaneous optical and classical electrophysiology. **A,** Simultaneously recorded *in vivo* optical (ArcLight) and electrical trace (interpolated at 2 kHz) from R5 neuron indicated in the wide-field imaging. **B,** Power spectrum analysis of membrane potential oscillations of R5 neuron shown in A. **C,** Cross correlograms indicating oscillatory character and temporal overlap of optical and electrical traces shown in A. **D,** Simultaneously recorded *in vivo* single-cell patch clamp (interpolated at 2 kHz) and optical compound activity of R5 dendritic area indicated in the wide-field imaging. **E,** Power spectrum analysis showing more complex rhythms in the optical compound activity. **F,** Amplitude cross correlation indicating partial temporal overlap between single-cell and compound activity.

Another important application for simultaneous patch clamp and voltage imaging is to investigate single-cell contributions to population dynamics reflected in compound recordings of neural activity. Here, we use NOSA to compare the activity of a single R5 ring neuron with the dorsal bulb, which is comprised of the dendrites of 10-12 R5 neurons (**Fig. 5D**). Power spectrum analysis shows that the peak frequency of the recorded R5 neuron is also represented in the dendritic compound signal (**Fig. 5E**). However, the power spectrum of the compound signal is much more complex due to the fact that the electrical patterns of several R5 neurons contribute to the compound signal. Correlation analysis suggests that some depolarized states of the single R5 neuron overlap with depolarized states in the compound signal (**Fig. 5F, compare Fig. 5D**).

### Cross correlation for analyzing event-based temporal relations

When imaging electrical activity in larger brain areas or even in whole brains NOSA can easily analyze the temporal relations of specific events occurring between different neuronal populations. To demonstrate this, we analyzed *in vivo* whole brain voltage imaging recordings in *Drosophila* (Aimon et al., 2019)(**Fig. 6A**). Stimulation with UV light induced electrical activity in the optic lobes while olfactory stimulation induced activity around the area of the lateral horns and peduncles of the mushroom bodies (**Fig. 6B**), which are both higher olfactory integration centers of the *Drosophila* brain (Heisenberg, 2003; Frechter et al., 2019). Interestingly, the central complex, which processes various sensory modalities (Green et al., 2017; Sun et al., 2017) and is important for basic locomotion (Strauss and Heisenberg, 1993; Strauss, 2002), shows spontaneous electrical activity. However, it is not clear whether some of the spontaneous activity originates from the visual or olfactory stimulation (**Fig. 6B**). Rather than detecting actual spikes, NOSAs’ spike detection can be used to determine the temporal relation between detected events. Events can be detected using a linear or dynamic threshold (**Fig. 1F**). Moreover, the threshold can be set either manually or based on the standard deviation of the whole recording. The detected events are used to generate a time-dependent event marker (**Fig. 6C, compare Suppl. Fig. 4**). The spike cross correlation function identifies the temporal relation between detected events, demonstrating that olfactory responses in the mushroom bodies and lateral horns are temporally aligned (**Fig. 6D**). Moreover, the set event markers and cross correlation analysis reveals that olfactory stimulation generates transient activity in the central complex, while visual stimulation does not lead to a detectable response (**Fig. 6D**).

**Figure 6.**
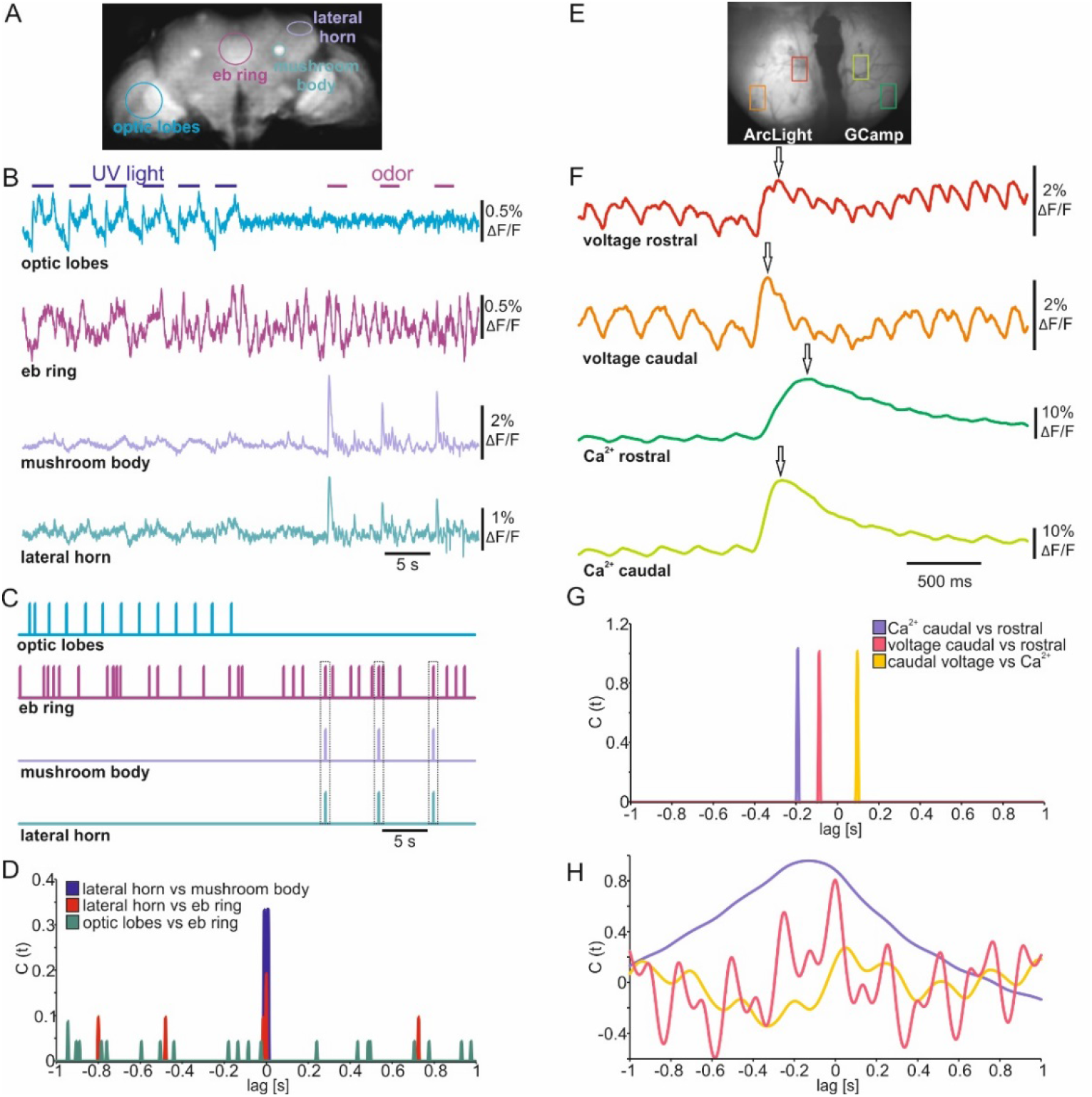
Event cross correlation identifies temporal relations in sensory evoked population dynamics. **A,** Wide-field image of a mid-section during whole brain voltage imaging (ArcLight) in *Drosophila*. **B,** Electrical population activity of different neuropils during optical and olfactory stimulation. **C,** Spike detection indicating detected events of electrical activity shown in B. **D,** Spike cross correlation indicating temporal shifts between events shown in C. Dashed squares indicate coincidence between olfactory evoked activity and specific events detected in the central complex. **E,** Wide-field image of the olfactory bulb injected with ArcLight and GCamp6f in either hemisphere. **F,** Olfactory evoked population activity of selected caudal and rostral glomeruli. Arrows indicate detected maxima. **G**, Spike cross correlation indicating temporal shifts between maxima indicated in F. **H**, Amplitude cross correlation indicating oscillatory character and temporal relations of population dynamics shown in F.

To analyze population dynamics in a mammalian brain we used a previously published recording in which one hemisphere of the olfactory bulb expresses ArcLight while the other expresses GCaMP6f (Storace et al., 2015) (**Fig. 6E**). The dorsal bulb exhibits a distinct temporal organization whereby glomeruli in the caudal bulb tend to be more coupled to respiration than glomeruli in the rostral olfactory bulb (Spors et al., 2006; Wachowiak et al., 2013). To demonstrate how quickly temporal relations can be established with NOSA we used the maxima of the olfactory responses in rostral and caudal glomeruli (**Fig. 6F**) to generate an event-based cross correlogram (**Fig. 6G**). This shows that electrical activity requires about 84 ms to travel from the caudal side of the olfactory bulb to the rostral side (**Fig. 6G**). In comparison, intracellular Ca^2+^ requires about 188 ms. Moreover, at the caudal glomeruli the delay between maxima of electrical activity and intracellular Ca^2+^ is 100 ms (**Fig. 6G**). In contrast to this event-based cross correlation the amplitude cross correlation (**Fig. 6H**) considers the whole recording and is thus influenced by the rhythmic changes in voltage which are a result of the mice’s breathing pattern (Storace et al., 2015).

## Discussion

Here, we report a novel open-source tool box, designed specifically for the analysis and interpretation of multicellular optical electrophysiology. While there is sophisticated software for processing imaging recordings (Romano et al., 2017), NOSA is entirely open access and, to our knowledge, the first analysis software that accounts for challenges specific to optical electrophysiology. In this manuscript we discuss these challenges and provide hands-on solutions to extract and analyze electrical patterns from recordings with low signal-to-noise ratio (SNR).

We demonstrate how high recording speeds necessary to resolve single action potentials drastically reduce the SNR (**Fig. 2, 3**). We therefore implemented baseline fitting and filtering algorithms, which can efficiently extract single action potentials and bursts from optical recordings (**Fig. 1-3**). During voltage imaging, the issue of a low SNR is aggravated by light scattering and movement artefacts for which we implemented background subtraction and movement correction algorithms (**Fig. 4**). We combine these basic but essential features for processing imaging data with sophisticated analysis tools for identifying electrical characteristics (**Fig. 3**) and functional interactions (**Fig. 2 and 6**). NOSA can also be used to investigate the temporal relation of sensory-evoked population dynamics in whole brain recordings in *Drosophila* (**Fig. 6A-D**) and in the olfactory bulb in mice (**Fig. 6E-H**). We here show that NOSA can be used to quickly identify temporal relations between the activity patterns of single cells and neuronal populations, which is crucial for investigating under which conditions neural networks interact with each other.

The properties and limitations of GEVIs affect the optical representation of neuronal activity. For example, the improved kinetics of the red-shifted GEVI Varnam increases the likelihood of resolving single spikes while ArcLight generally yields higher SNR (**Fig. 2**). NOSA can be used to quickly determine the properties of GEVIs. Using NOSA we verify that the GEVI Ace2N has faster kinetics than ArcLight (Gong et al., 2015), especially with respect to the repolarization phase of an action potential (**Fig. 3B**). However, the temporal representation of bursts is similar in both GEVIs (**Fig. 3G**). In fact, during bursts ArcLight yields higher relative changes in fluorescence (**Fig. 3G**). With specific knowledge about the properties and limitations, the adequate GEVI can be chosen for a specific experiment. A comprehensive characterization of the properties of different GEVIs are reported elsewhere (Bando et al., 2019).

NOSA’s event detection can be used to extract neuron-specific firing characteristics (**Fig. 3**), enabling the fast identification of specific types of neurons within a population of seemingly homogeneous neurons. This knowledge could then be used to electrically stimulate neurons with specific attributes and analyze their connectivity to other neurons (Antic et al., 2016). Such sophisticated experiments would benefit from another feature, which is provided by NOSA: the simultaneous analysis of optical and electrical data (**Fig. 5**). Moreover, we show how this feature can be used to analyze single-cell contributions to population dynamics (**Fig. 5D**). This is especially important when trying to understand how a multitude of neurons orchestrate their electrical activities to generate population dynamics, e.g. during sleep (Buzsaki and Draguhn, 2004).

GEVIs are currently improving at a rapid pace, developing towards stronger fluorescence and improved kinetics. However, currently the diverse properties of GEVIs and a missing analytical pipeline represent motivational bottlenecks preventing experimental implementation and widespread use of GEVIs. With NOSA we provide an analytical toolbox that will greatly facilitate the use of GEVIs in studying multicellular electrical patterns, inexorably improving our understanding of functional interactions within neural networks.

## Supplementary Figures

**Supplementary figure 1:**
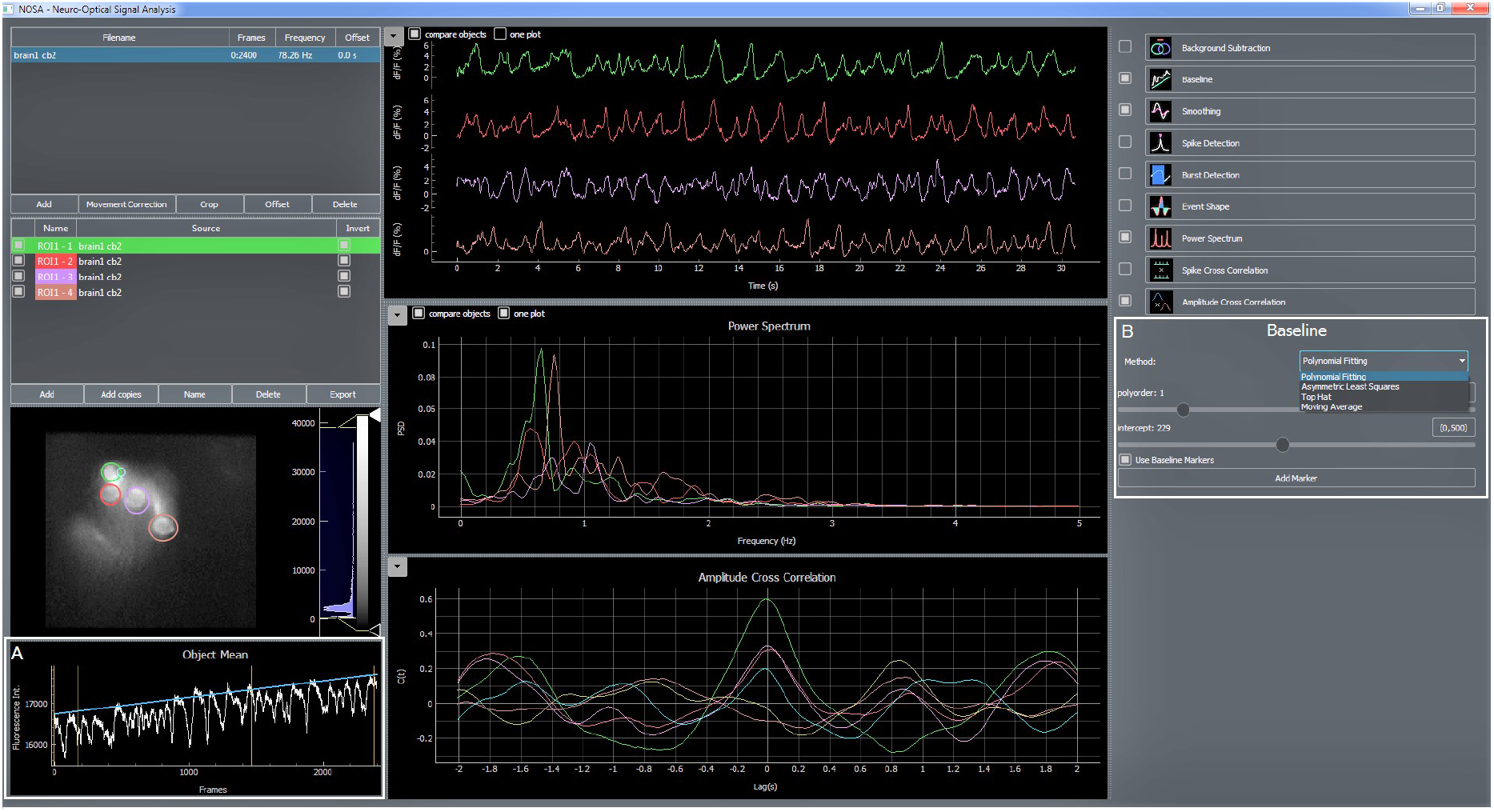
NOSA baseline correction. Related to Figure 2. **A,** Polynomial fitting (blue curve) is used to compensate for baseline drifts and calculate the relative change in fluorescence. Additional baseline markers (orange) can be used to guide the fitting curve. **B**, NOSA provides four different algorithms for baseline fitting. For each algorithm, the baseline can be adjusted by changing the shape (polyorder) and position (intercept) of the fitting curve.

**Supplementary figure 2:**
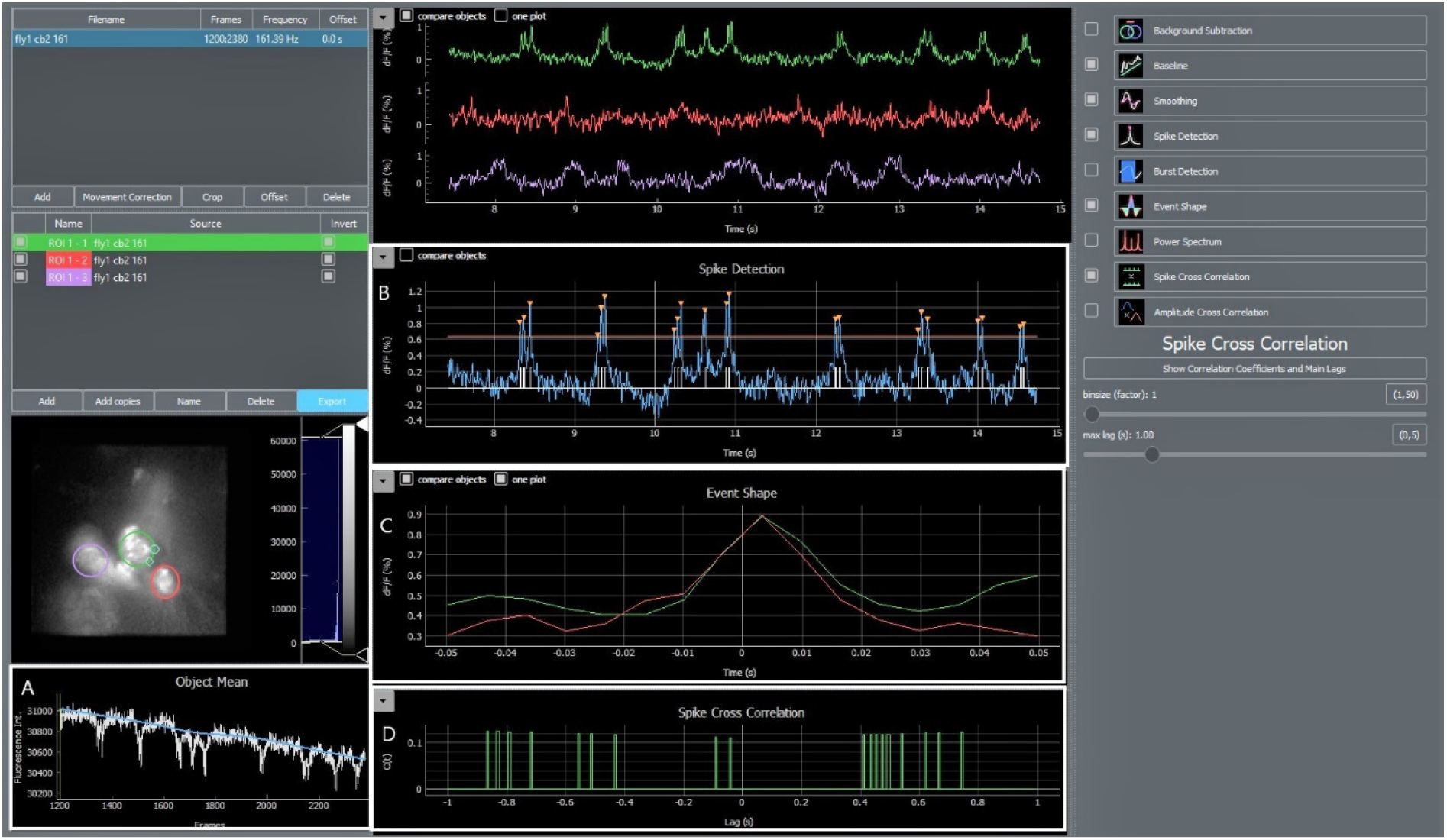
NOSA spike detection and cross correlation. Related to Figure 2. **A,** Shift in baseline fluorescence is corrected via the asymmetric least square algorithm (blue fitting curve). **B,** Spike detection using a linear threshold approach. Orange arrows indicate detected spikes. **C,** Event shape can be used to generate the average shape of spikes detected in B. **D,** NOSA spike cross correlation can be used to analyze the temporal relation between spikes detected in cell 1 (green) and cell 2 (red).

**Supplementary figure 3:**
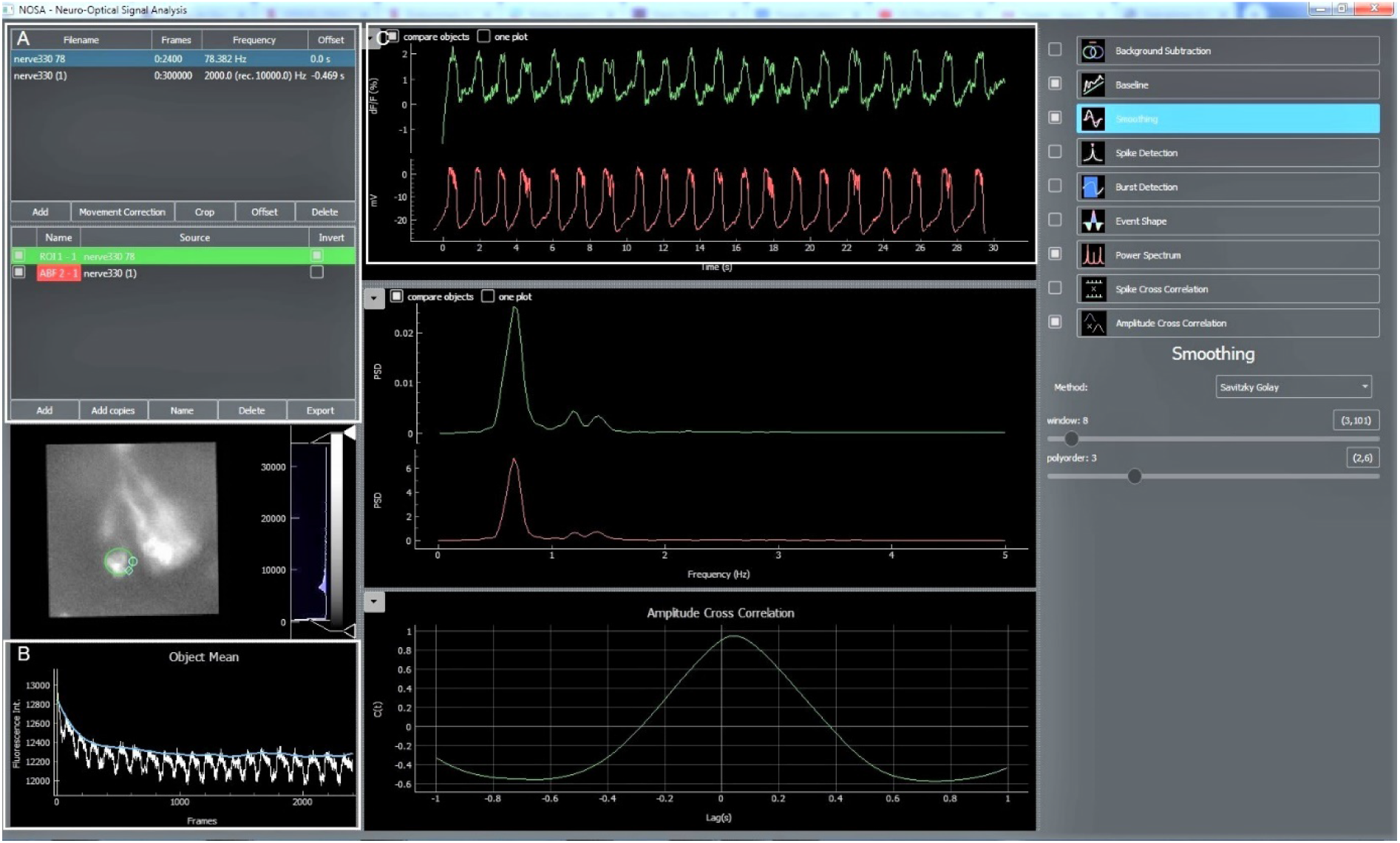
Simultaneous analysis of optical and electrical recordings. Related to Figure 5. **A**, Optical (tiff files) and electrical (abf files) recordings can both be imported into NOSA. Sampling rates and recording speeds can be adjusted via various interpolation algorithms. The sampling rate of the electrical recording (10 kHz) was adjusted to 2 kHz. Delays between the electrical and optical systems can be corrected by an offset function (see C). **B**, Optical recording was corrected for bleaching via the asymmetric least square algorithm. **C**, The offset function in NOSA (see A) can adjust for temporal delays between the electrical and optical recordings. In this case, the precise delay was 469 ms.

**Supplementary figure 4:**
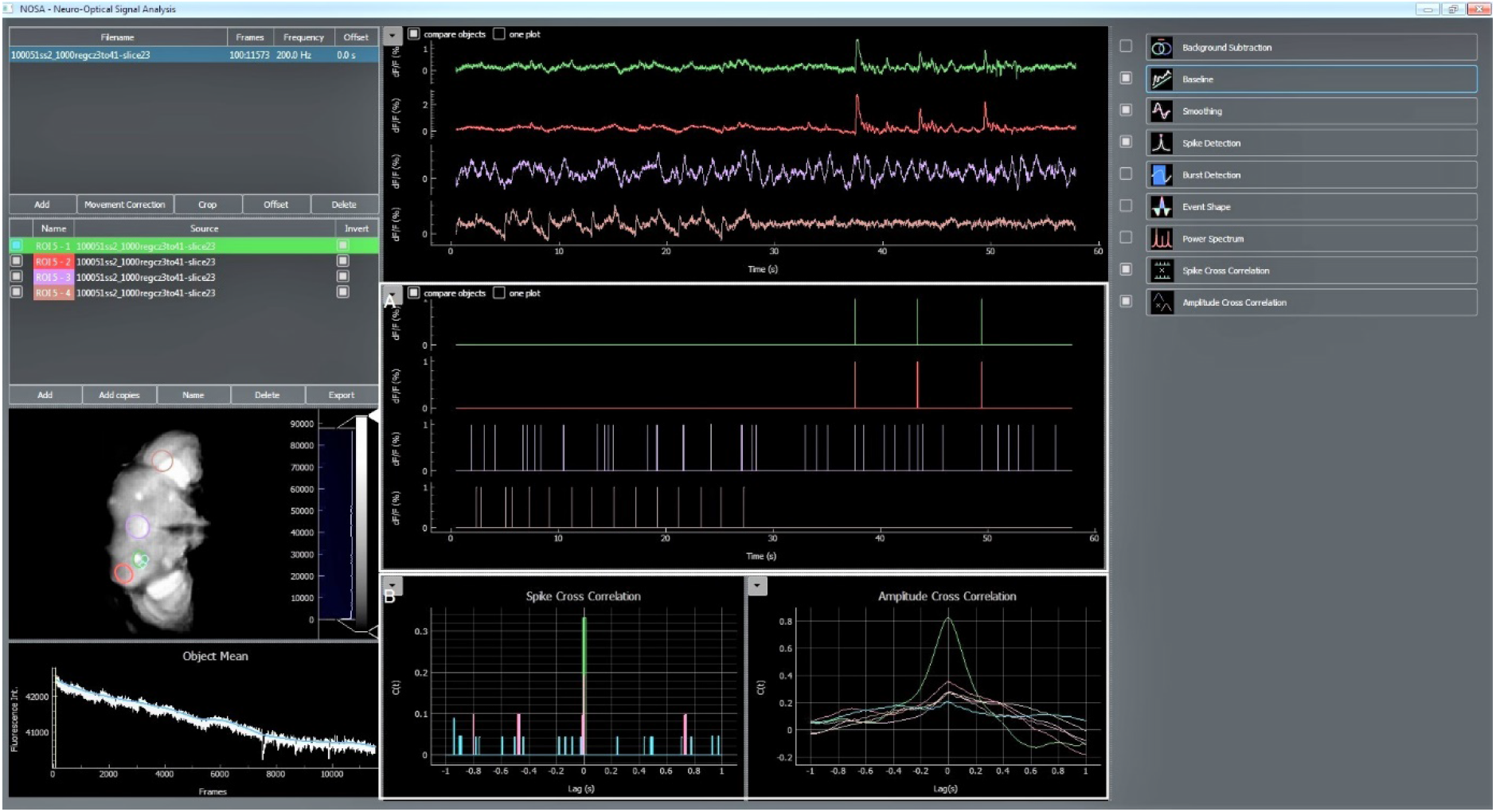
Combining event detection and cross correlation to identify temporal relations in population dynamics. Related to Figure 6. **A**, Spike detection can be used to detect specific events, generating a time-dependent event marker. **B**, Spike cross correlation calculates the temporal relation between detected events shown in A. In comparison, the amplitude cross correlation calculates temporal relations based on the relative change in fluorescence.

## Methods

### NOSA software

NOSA was written in <u>Python</u> 3.7.1 (see https://docs.python.org/3/license.html for license information). Besides default packages and built in functions, NOSA uses a variety of additional packages (**Table 1**). More detailed information on the graphical user interface, the function of specific features, performance aspects and workflow examples are provided in an additional file accompanying the software file. For more information please send an email to the lead contact.

**Table 1:**
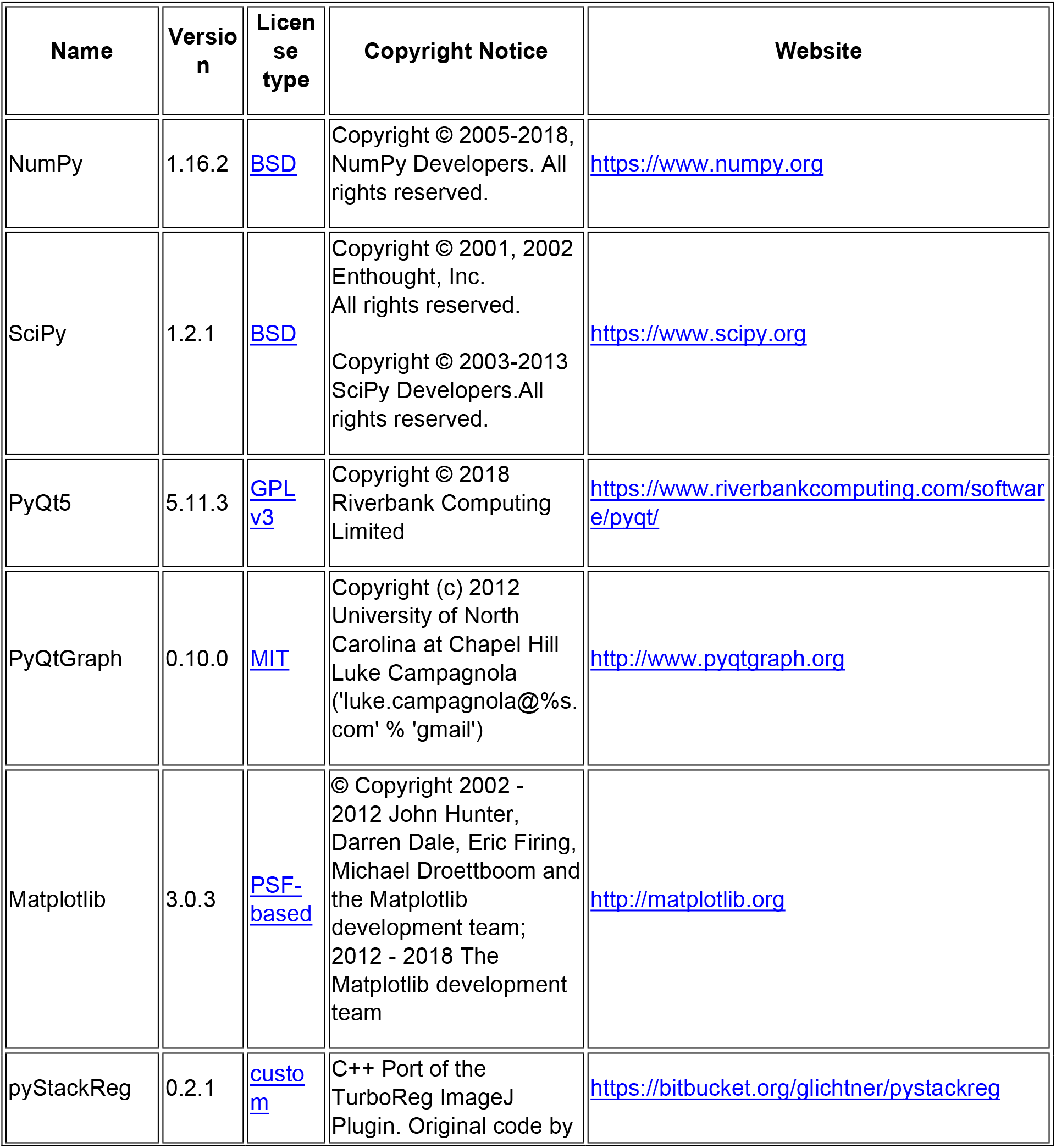
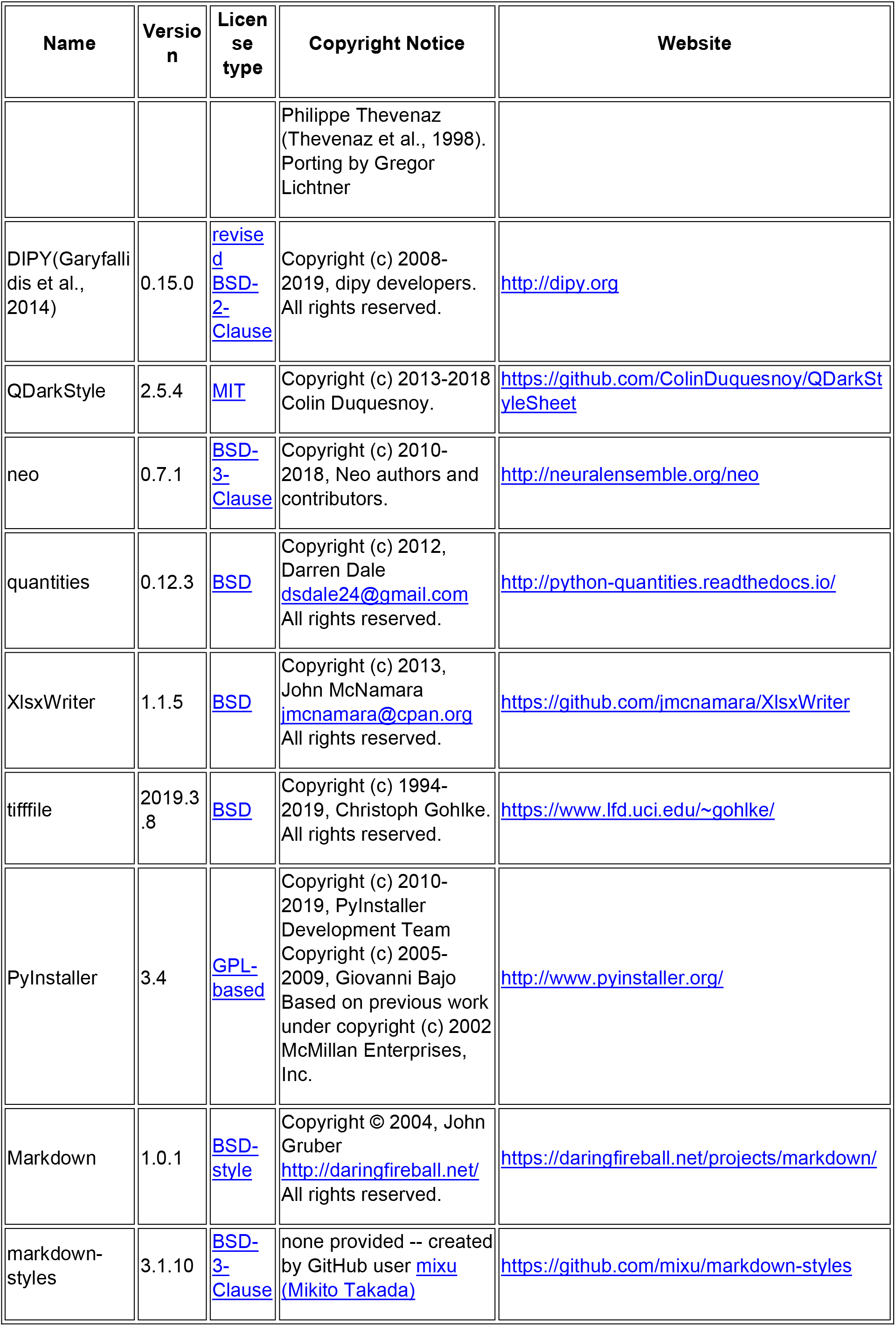
Packages used for NOSA

### Optical and classical electrophysiology of R5 neurons in Drosophila melanogaster

Flies (*Drosophila melanogaster*) were reared on standard cornmeal food at 25 °C and 60% humidity under a 12 h light/dark regime. All flies were obtained from the Bloomington Stock resource center (BDRC). Experiments were performed with 3-10 d old female flies at ZT 8-12 (Zeitgeber time; during a 12/12 h light/dark cycle the onset of light is at ZT 0 h and the offset of light is at ZT 12 h). Whole-brain explant dissections and fly *in vivo* preparation were performed as previously described (Cao et al., 2013; Owald et al., 2015). During ex vivo experiments (**Fig. 2, 3**) external solution consisted of (in mM): 90 NaCl, 3 KCl, 1.5 CaCl_2_, 5 MgCl_2_, 1 NaH_2_PO_4_, 10 glucose, 10 sucrose, 8 trehalose, 5 TES and 26 NaHCO_3_. During ex vivo experiments (**Fig. 4, 5**) external solution consisted of (in mM): 70 NaCl, 3 KCl, 1.5 CaCl_2_, 20 MgCl_2_, 1 NaH_2_PO_4_, 10 glucose, 10 sucrose, 8 trehalose, 5 TES and 26 NaHCO_3_.

Imaging was performed on an Olympus BX51WI microscope using a Plan Apochromat 40×, numerical aperture 0.8, water-immersion objective (Olympus, Japan). ArcLight was excited at 470 nm using a Lumencor Spectra X-Light engine LED system. LED power was adjusted for each recording individually to make sure that fluorescent images were not saturated. The objective C-mount image was projected onto an Andor iXon-888 camera controlled by Andor Solis software. Imaging was performed at frame rates of 80 Hz (**Fig. 2A-C, 4 and 5**), 160 Hz (**Fig. 2D-G**) 250 Hz and 1000 Hz (**Fig. 3**).

*In vivo* (**Fig. 5**) whole-cell patch-clamp recordings from R5 neurons were performed at ZT 8-12 as reported elsewhere (Wilson and Laurent, 2005; Murthy and Turner, 2013; Donlea et al., 2018). Neurons were recorded for up to 5 min. Identification of R5 neurons was based on ArcLight expression. External saline was used as described above. Patch pipettes (7-10 MΩ) were filled with internal saline containing (mM): 135 K-aspartate, 10 HEPES, 1 EGTA, 1 KCl, 4 MgATP, 0.5 Na3GTP. Internal solution was adjusted to a pH of 7.2, with an osmolarity of 265 mmol/kg.

### Ca^2+^ imaging of hippocampal slices

Animal care and handling was in accordance with the Helsinki declaration and institutional guidelines. Protocols for organ removal and culturing were approved by the State Office of Health and Social Affairs Berlin, under the license number T0123/11. Organotypic hippocampal slice cultures were prepared as described previously (Kann et al., 2011; Prager et al., 2019). In short, hippocampal slices (400 μm) were obtained from Wistar rat pups at postnatal day 6-7. Slices were positioned on cell culture membrane inserts (Millicell-CM, Millipore) and maintained for 6 days in culture medium (50% MEM, 25% HBSS, 25% Horse Serum and 1 mM L-glutamine, pH set to 7.3) at 5% CO_2_. At day 7, slice cultures were bulk stained with OGB-1-AM BAPTA (5μM in DMSO, 0.01% Pluronic F-127, 0.005% Cremaphor) by submerging them for 50 min at room temperature in carbogen bubbled serum free medium.

Fluorescence recordings were obtained with a spinning disk confocal microscope (Andor Revolution, BFIOptilas GmbH, Gröbenzell, Germany), equipped with an EMCCD camera (Andor iXonEM+, 60x objective N.A.1, 2 x binning, 20 Hz recording speed, laser intensity 150-200 μW at the focal level). Slice cultures were superfused with carbogen (95% O_2_, 5% CO_2_) saturated artificial cerebrospinal fluid containing (in mM): 129 NaCl, 3 KCl, 1.25 NaH_2_PO_4_, 1.8 MgSO_4_, 1.6 CaCl_2_, 26 NaHCO_3_ and 10 glucose (pH 7.4, t=30°C). For induction of synchronized epileptiform activity, Mg^2+^ was omitted and KCl was elevated to 5 mM (Prager et al., 2019).

### Whole brain voltage imaging in Drosophila

Whole brain recordings were performed using light field microscopy as described in detail elsewhere (Aimon et al., 2019). In short, a modified upright Olympus BX51W with a 20x NA 1.0 XLUMPlanFL (Olympus) was used. An adequate microlens array (RPC Photonics) positioned at the image plane and two relay lenses (50 mm f/1.4 NIKKOR-S Auto from Nikon) projected the image onto the sensor of a scientific CMOS camera (Hamamatsu ORCA-Flash 4.0). A 490 nm LED (pE100 CoolLED) at approximately 10% of its full power was used for excitation. As filter set we used a 482/25 bandpass filter, a 495-nm dichroic beam splitter, and a 520/35 bandpass emission filter (BrightLine, Semrock). The recording was performed at a frame rate of 200 Hz. The whole brain volume was reconstruction from the light field image as described in (Aimon et al., 2019)

### Voltage and calcium imaging in the olfactory bulb

Olfactory bulb recordings in mice were performed as described in detail elsewhere (Storace et al., 2015). In short, C57BL/6 mice (JAX, Bar Harbor, MA) were injected into the olfactory bulbs with AAV1 expressing either ArcLight- or GCaMP6f. Mice were anesthetized (ketamine/xylazine) and the bone above both olfactory bulbs was either thinned or removed. The exposure was covered with agarose and sealed with a glass coverslip. The dorsal surface of both hemispheres was illuminated with 485 ± 25 nm light using epifluoresence illumination on a Leitz Ortholux II microscope with a tungsten halogen lamp or a 150 W Xenon arc lamp (Opti Quip) and a 515 nm long-pass dichroic mirror. Fluorescence emission was recorded with a NeuroCCD-SM256 camera with 2 × 2 binning at 125 Hz using NeuroPlex software (RedShirtImaging, Decatur, GA). All surgical procedures were approved by the Yale IACUC.

